# Overdominant mutations restrict adaptive loss of heterozygosity at linked loci

**DOI:** 10.1101/2021.05.13.444075

**Authors:** Kaitlin J. Fisher, Ryan C. Vignogna, Gregory I. Lang

## Abstract

Loss of heterozygosity is a common mode of adaptation in asexual diploid populations. Because mitotic recombination frequently extends the full length of a chromosome arm, the selective benefit of loss of heterozygosity may be constrained by linked heterozygous mutations. In a previous laboratory evolution experiment with diploid yeast, we frequently observed homozygous mutations in the *WHI2* gene on the right arm of Chromosome XV. However, when heterozygous mutations arose in the *STE4* gene, another common target on Chromosome XV, loss of heterozygosity at *WHI2* was not observed. Here we show that mutations at *WHI2* are partially dominant and that mutations at *STE4* are overdominant. We test whether beneficial heterozygous mutations at these two loci interfere with one another by measuring loss of heterozygosity at *WHI2* over 1,000 generations for ∼300 populations that differed initially only at *STE4* and *WHI2*. We show that the presence of an overdominant mutation in *STE4* reduces, but does not eliminate, loss of heterozygosity at *WHI2*. By sequencing 40 evolved clones, we show that populations with linked overdominant and partially dominant mutations show less parallelism at the gene level, more varied evolutionary outcomes, and increased rates of aneuploidy. Our results show that the degree of dominance and the phasing of heterozygous beneficial mutations can constrain loss of heterozygosity along a chromosome arm, and that conflicts between partially dominant and overdominant mutations can affect evolutionary outcomes.

**SIGNIFICANCE STATEMENT:** In diploid populations, it is beneficial for partially dominant beneficial mutations to lose heterozygosity, but it is deleterious for overdominant beneficial mutations to do so. Because loss-of-heterozygosity tracts often encompass entire chromosome arms, a conflict will arise when a partially dominant beneficial mutation and an overdominant beneficial mutation exist in close proximity. We demonstrate that this conflict occurs, and that it restricts loss of heterozygosity, resulting in more variable evolutionary outcomes.

## INTRODUCTION

The pace of adaptation in asexual diploids is strongly dependent on the dominance of new beneficial mutations. Theoretically, the probability of a given beneficial mutation fixing in a population is the product of its coefficient of selection (*s*) and its degree of dominance (*h*), where *h* = 0 is fully recessive and *h* = 1 is fully dominant (Orr and Otto 1994). Beneficial alleles with a low degree of dominance (*h* ≈ 0) are unlikely to fix in asexual diploid populations, a phenomenon known as Haldane’s Sieve (Haldane 1924, Connallon and Hall 2018). Depending on the degree of dominance, some adaptive pathways that are open to haploids, will be improbable or inaccessible to diploids. Changes to the spectrum of beneficial mutations and to the genetic targets of selection between haploid and diploid populations provide experimental evidence of this constraint (Fisher *et al*. 2018, Marad *et al*. 2018).

Recessive beneficial mutations can be converted to beneficial homozygous mutations through loss-of-heterozygosity (LOH) events during asexual propagation of diploid yeast populations (Gerstein *et al*. 2014). For highly heterozygous populations, such as inter-specific hybrids, LOH becomes the dominant mechanism of adaptation (Smukowski Heil *et al*. 2017, James *et al*. 2019) due to a high rate of LOH relative to point mutation (Barbera and Petes 2006, Lee *et al*. 2009) and to a reservoir of beneficial mutations that are masked in the heterozygous state. The ability of a given allele to escape Haldane’s sieve by LOH will depend on its genomic location since the rate of LOH varies across the yeast genome (Lee *et al*. 2009). In natural isolates the rate of LOH increases with distance from the centromere (Peter *et al*. 2018) and in experimental evolution several hotspots for LOH have been identified, most strikingly at the rDNA locus on Chromosome XII (Fisher *et al*. 2018, Marad *et al*. 2018).

Two special cases of dominance further constrain adaptation in diploids. Underdominance (*h* < 0) occurs when the heterozygote is less fit than either homozygous genotype. Underdominant mutations are unlikely to establish as heterozygotes and therefore underdominance impedes access to potentially adaptive homozygous genotypes. At the other extreme, overdominance (*h* > 0) occurs when the heterozygote is more fit than either homozygous genotype. Overdominant mutations should readily establish in populations and be maintained as heterozygous by selection against homozygous genotypes (Fisher 1928). Because little is known about the distribution of the degree of dominance of new mutations in diploids, the relative importance of underdominant and overdominant mutations in genome evolution is unclear. Theoretically, overdominant mutations are predicted to be a major contributor to the maintenance of genetic variation (Maruyama and Nei 1981), and a frequent, if perhaps only transient, outcome of diploid evolution (Manna *et al*. 2011, Sellis *et al*. 2011). Though genome-wide scans have turned up little evidence of overdominance (Szulkin *et al*. 2010, Hedrick 2012, Goudie *et al*. 2014), laboratory-evolution experiments demonstrate that overdominant mutations can contribute to short-term adaptation in diploid populations (Sellis *et al*. 2016, Leu *et al*. 2020).

If overdominant mutations are frequent in evolving asexual diploid populations, fitness conflicts may arise when overdominant (*h* > 0) and partially dominant (0 < *h* < 1) heterozygous beneficial mutations arise in close proximity to one another on a chromosome. This is because an LOH event would convert both mutations to the homozygous state resulting in a fitness loss due to the overdominant mutation and a fitness gain due to the partially dominant mutation. We demonstrate that this conflict arises in experimental evolution between an overdominant mutation in *STE4* and a partially dominant mutation in *WHI2*, both of which are located on the right arm of Chromosome XV. We show experimentally that adaptive LOH at the *WHI2* locus is slowed by the presence of the overdominant *STE4* mutations.

## RESULTS

We previously identified 20 genes that are targets of selection in 46 laboratory-evolved populations yeast that were propagated asexually for 4,000-generations (Fisher *et al*. 2018). By generation 1,000, all 46 populations had autodiploidized and therefore most adaptation occurred in the diploid state. Unlike true diploids (*MAT***a**/α), which do not mate, autodiploids (*MAT***a**/**a** or *MAT*α/α) produce mating pheromones and will readily mate with cells of the opposite mating type. Therefore, autodiploids, like haploids (*MAT***a** or *MAT*α), should benefit from mutations that inactivate the mating pathway (Lang *et al*. 2009). However, only one mating pathway gene, *STE4*, is identified as a common target of selection across the 46 autodiploid populations (Fisher *et al*. 2018). In contrast, mutations in *STE4, STE5, STE11*, and *STE12* are overrepresented in 40 closely matched haploid populations (Lang *et al*. 2013). Though not identified in either experiment, *STE7* is also a target of selection in haploids (Lang *et al*. 2009, Buskirk *et al*. 2017). We have demonstrated previously that mating pathway mutations are loss-of-function based on a combination of their mutational spectra, allele reconstruction, and gene deletion (Lang *et al*. 2009, Lang *et al*. 2013, Buskirk *et al*. 2017). However, *STE4* mutations in autodiploids are inconsistent with simple loss-of-function: not only are they found in only one gene, but all six *STE4* mutations are heterozygous and cluster in a small (260 bp) region of the coding sequence (*X*^*2*^(1, *N=6*) = 18.76, *p*<10^−4^, **Figure 1A**).

**Figure 1.**
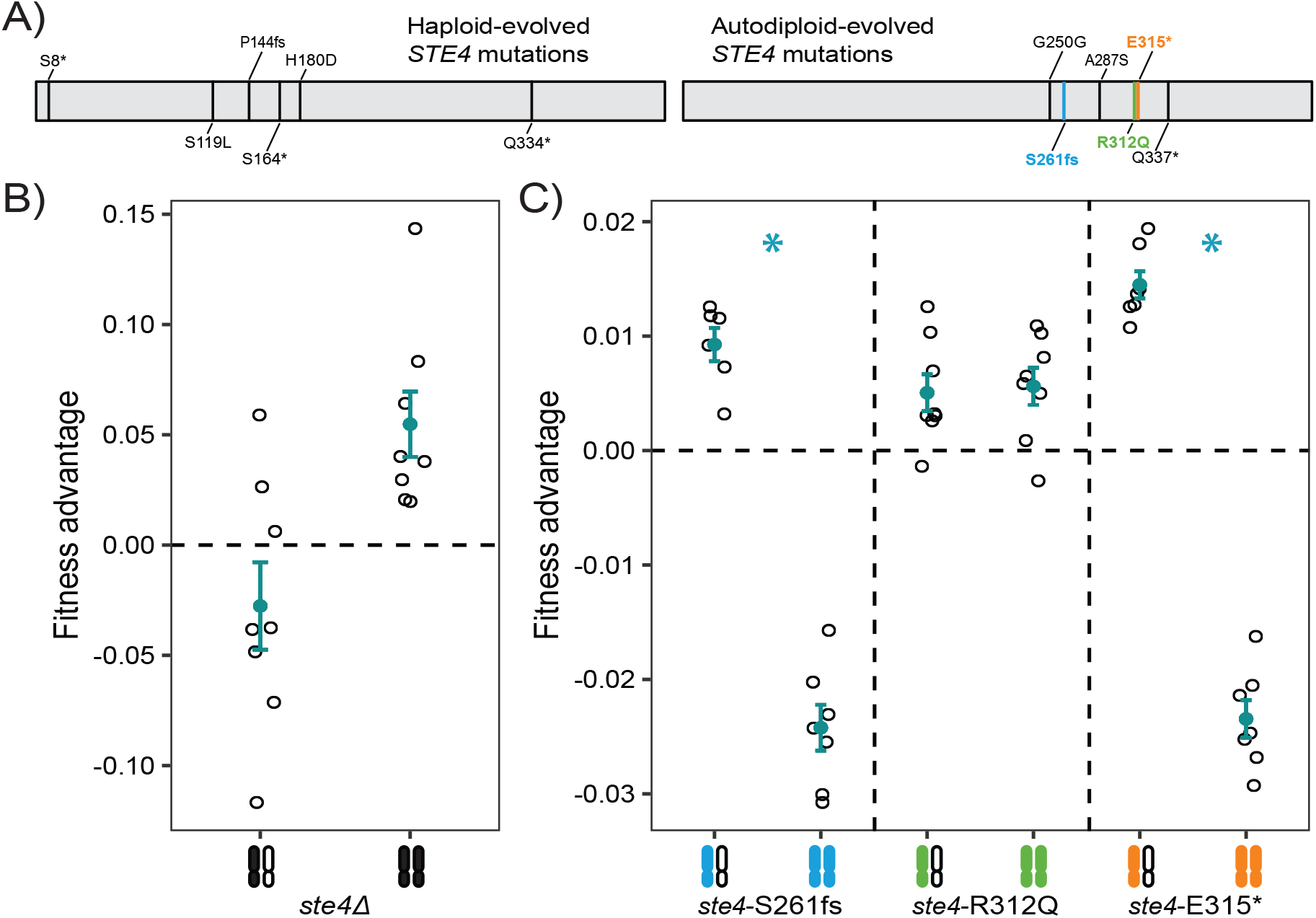
Adaptive *STE4* mutations differ between haploid and autodiploid populations. **A)** Mutations previously identified in the *STE4* gene in haploid (Lang *et al*. 2013) and autodiploid (Fisher *et al*. 2018) populations. The positions of mutations across the coding sequence of *STE4* in haploid populations did not deviate from random expectation (*X*^*2*^(1, *N=6*) = 0.74363, *p*=0.39). Conversely, autodiploid mutations accumulated nonrandomly across the linear 1,276 bp sequence of the *STE4* gene, (*X*^*2*^(1, *N=6*) = 18.76, *p*<10^−4^). Bolded mutations are ones that were selected for reconstruction. **B)** Deletion of *STE4* is deleterious in autodiploids when heterozygous and beneficial when homozygous. **C)** Three evolved autodiploid alleles were reconstructed in an ancestral background. *ste4*-S261fs and *ste4*-E315* are overdominant in the ancestral background. **B-C**) A filled and open chromosome represents heterozygosity and two filled chromosomes represents homozygosity. Open points are fitness measures of eight biological replicates following *MAT***a**/**a** conversion. Filled points show mean fitness **±** standard error.

*STE4* encodes the highly conserved beta subunit of the heterotrimeric G protein complex. We mapped the positions of the evolved mutations onto the homology-predicted structure of Ste4p (**Figure S1)**. All six autodiploid mutations impact residues near the C-terminal end of the protein, with three of these (a frameshift and two nonsense mutations) resulting in truncations of the final ∼100 amino acids. Two missense mutations both occur in a putative random coil that derives from a yeast-specific insertion (Sondek *et al*. 1996). One synonymous mutation also arose in this region.

### Adaptive *STE4* mutations in autodiploids are either dominant or overdominant

We hypothesized that the discrepancy between patterns of sequence evolution in haploids and autodiploids is because the beneficial mating pathway mutations in haploids are recessive. We first tested whether the fitness benefit of evolved *ste4* mutations in autodiploids is phenocopied by gene deletion, as is the case with *ste4* mutations arising in haploid populations. We generated *STE4* deletion (*ste4*Δ) strains as haploids, as heterozygous autodiploids, and as homozygous autodiploids. As reported previously, we find that *ste4*Δ mutants are beneficial in a haploid background (**Figure S2)**. Similarly, homozygous *ste4*Δ/*ste4*Δ mutants are beneficial in *MAT***a**/**a** diploids (**Figure 1B**). Surprisingly, however, heterozygous *STE4*/*ste4*Δ mutants are underdominant: substantially less fit than the either homozygous genotype (**Figure 1C**).

The underdominance of *STE4* deletions confirms that evolved autodiploid mutations are not purely loss-of-function, as these would be deleterious. We used CRISPR-Cas9 allele-swaps to reconstruct three autodiploid-evolved *ste4* alleles (one frameshift, one nonsense, and one missense mutation, **Figure 1A**). For each allele, we then assayed the fitness of the haploid mutant and the homozygous and heterozygous autodiploid mutants. In a haploid background the evolved frameshift and nonsense alleles have a fitness benefit (2.1 ± 0.3% and 2.2 ± 0.4%, respectively; mean ± standard error, *p*<10^−4^ both genotypes) while the missense allele is neutral (−0.7 ± 0.3%, *p*=0.995, **Figure S2**; Note that we also verified that the synonymous PAM site mutation is neutral, **Figure S3**).

In the heterozygous autodiploids, like in the haploids, the *ste4*-S261fs and *ste4*-E315* mutations are beneficial (0.8 ± 0.2% and 1.2 ± 0.1%, *p=*0.02 and *p*<10^−4^, respectively), and the *ste4*-Arg312Gln mutation is neutral (0.2 ± 0.2%, *p*=0.49, **Figure 1C**). The fitness effect of the heterozygous evolved alleles is ∼40-50% of the fitness advantage conferred in haploids. Given that 95% of mutations in the autodiploids are heterozygous after 4,000 generations it is unsurprising that all six *STE4* mutations are heterozygous, and we expected that the homozygous mutations would be equally (if not more) fit than the heterozygous *STE4* mutations. However, the *ste4*-S261fs and *ste4*-E315* homozygous mutants have fitness defects of −2.1 ± 0.1% and −2.4 ± 0.1%, respectively (mean ± standard error, *p*<10^−3^ for both comparisons, **Figure 1C**). Therefore, rather than showing an additive fitness effects as predicted, two *STE4* missense mutations are overdominant (*h* > 1).

### Linkage to overdominant *STE4* alleles delays, but does not prevent, adaptive LOH

We next examined whether, in our populations, the presence of overdominant *ste4* mutations constrains adaptive LOH at linked loci. Although most mutations in our evolved autodiploids are heterozygous, there are two large genomic regions that are prone to high rates of loss-of-heterozygosity (LOH); these regions are identifiable based on the clustering of homozygous mutations in evolved genomes (Fisher *et al*. 2018). One of these regions on the right arm of Chromosome XV contains the *STE4* locus as well as three other common targets of selection in our experimental system: *WHI2, SFL1*, and *PDR5*. Unlike *STE4*, putative adaptive mutations in *WHI2, SFL1*, and *PDR5* are commonly observed to be homozygous in evolved clones. We show that an evolved mutation in *WHI2* (*whi2*-Q29*) is partially dominant, having a 3.4 ± 0.2% benefit when heterozygous and a 5.1 ± 0.3% benefit when homozygous (mean ± standard error, **Figure 2A**).

**Figure 2.**
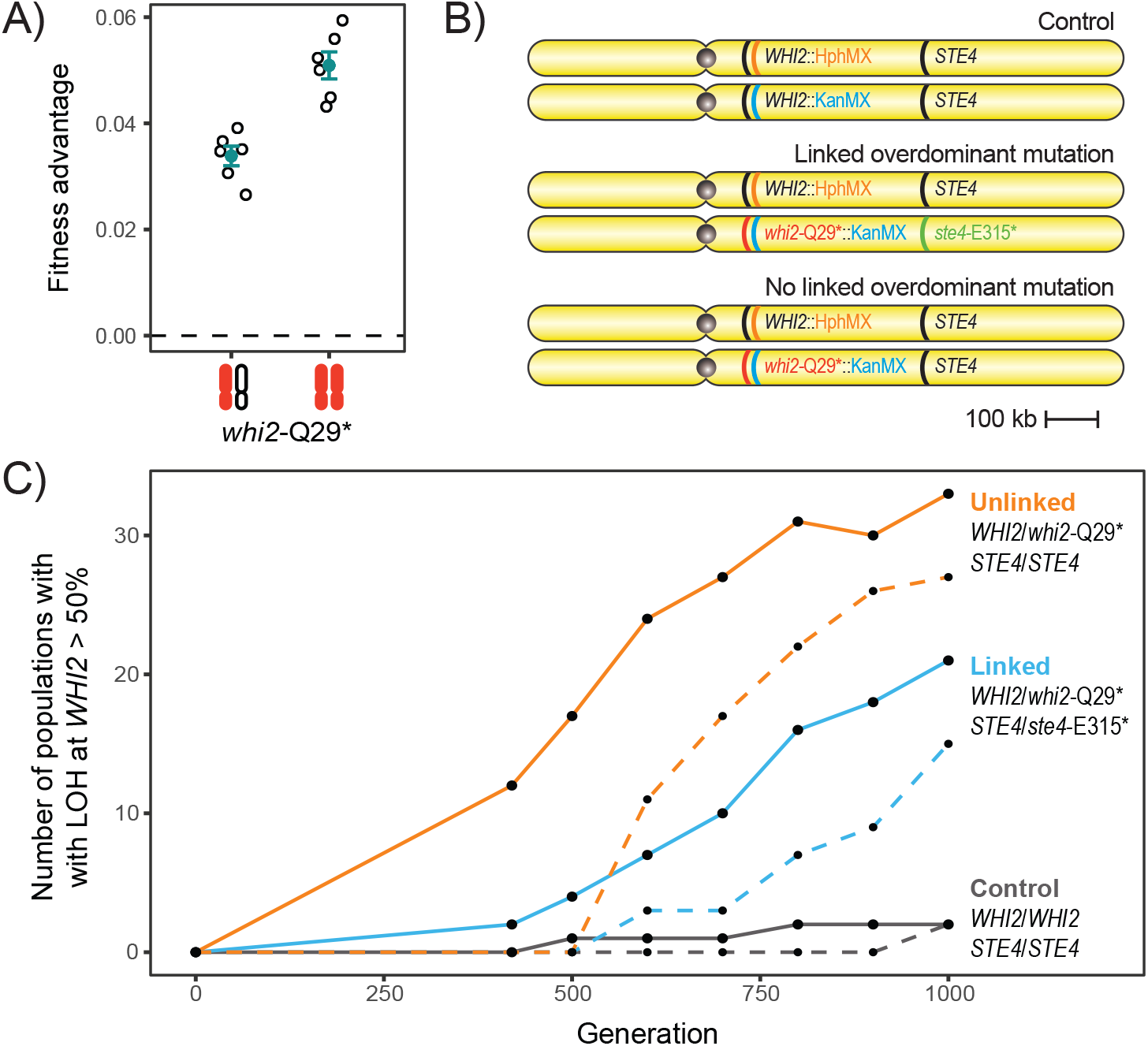
The presence of a linked overdominant *STE4* mutation restricts, but does not prevent, loss of heterozygosity at *WHI2*. **A)** An evolved *Q29\** in the *WHI2* gene, which is centromeric to *STE4*, reported in (Fisher *et al*. 2018) is partially dominant and most beneficial when homozygous. A filled and open chromosome represents heterozygosity and two filled chromosomes represents homozygosity. Open points are fitness measures of eight biological replicates following *MAT***a**/**a** conversion. Filled points show mean fitness **±** SE. **B)** Schematic of the three genotypes that were constructed for an evolution experiment. LOH in any genotype is determined by loss of double drug resistance and the direction of the LOH is determined by the single drug to which resistance is lost. **C)** 288 populations (96 *whi2*-Q29*/*WHI2 STE4*/*STE4*, 96 *whi2*-Q29*/*WHI2 ste4*-E315*/*STE4*, 95 wild-type control) were evolved for 1,000 generations. Solid lines show the number of populations with detected LOH over time. Dashed lines show the number of populations in which *whi2*-Q29*homozygous genotypes were fixed. Lines are colored by group (orange: *ste4*-E315* linked, blue: *STE4* wild-type, black: control). LOH of *whi2*-Q29* was observed in a higher fraction of unlinked populations at generation 420 (Fisher’s exact, *p*=0.001), however, the difference becomes smaller over time and is nonsignificant at generation 1,000 (Fisher’s exact, *p*=0.08). At generation 1,000, *WHI2* LOH was detectable in 33 *whi2*-Q29* *STE4* populations, 21 *whi2*-Q29* *ste4*-E315* populations, and 2 control populations.

*WHI2, SFL1*, and *PDR5* are all centromere proximal to *STE4*, and since conversion tracts produced by mitotic recombination frequently extend from a medial breakpoint to the telomere, adaptive LOH of any of these three loci is likely to extend to through the *STE4* locus. We examined evolved autodiploid genotypes for evidence of LOH events occurring after a *STE4* mutation arises on the right arm of Chromosome XV and we find none. There are two populations with fixed homozygous Chromosome XV mutations, but each contain heterozygous *STE4* alleles, indicating that the LOH event on Chromosome XV occurred before *STE4* mutations in these populations. The inverse—unfixed Chromosome XV homozygous mutations on a fixed *ste4* mutant background, which would indicate LOH on Chromosome XV after a mutation at *STE4*—is not observed in any of the three populations with fixed *STE4* mutations (**Figure S4**).

To explicitly test the hypothesis that overdominant *STE4* alleles decrease the likelihood of adaptive LOH at linked loci, we performed an evolution experiment using three strains that differed only on the right arm of Chromosome XV (**Figure 2B**). One strain contained a heterozygous beneficial and partially dominant *whi2*-Q29* mutation and was wild-type at the *STE4* locus (*WHI2*/*whi2*-Q29*, *STE4/STE4*). A second strain contained the same heterozygous *whi2*-Q29* mutation in *cis* with an a heterozygous overdominant *ste4*-E315* mutation (*WHI2*/*whi2*-Q29*, *STE4/ste4*-E315*). A control strain was wild type at both loci.

We evolved 96 replicate populations of each strain (95 for control strain) for 1,000 generations. We tested for LOH events at the *WHI2* locus every 100 generations starting at Generation 400. All three strains carried KanMX and HphMX drug-resistance cassettes tightly linked to the *WHI2* loci on each chromosome. By assaying for loss of the ability to grow on double drug (G418+Hygromycin), our assay is sensitive to LOH events that reach frequency of 0.5 in the population (**Figure S5**).

The presence of a heterozygous and partially dominant *whi2*-Q29* adaptive mutation drives a high rate of fixation of LOH events relative to the wild-type control populations (**Figure 2C**). However, among populations with a *whi2*-Q29* allele, the fraction of populations experiencing LOH is significantly lower in populations with a telomeric overdominant *ste4*-E315* mutation at Generation 420 (Fisher’s exact, *p*=0.001, **Figure 2C**) and remains lower through all time points assayed, although this becomes non-significant at Generations 900 and 1,000 (Fisher’s exact, *p*=0.066, 0.077). While time-course LOH dynamics show an effect of linkage to an overdominant allele, the linkage did not prevent LOH at *WHI2*, as would be predicted by the additive fitness effects of both homozygous mutations (**Figures 1C and 2B**).

Although both the *WHI2* and *STE4* loci are on the right arm of Chromosome XV, they are 700 kb apart. We hypothesized that LOH events that occurred in populations with both *WHI2*/*whi2*-Q29* and *STE4/ste4*-E315* heterozygosities might involve short tracts of mitotic recombination that included the *WHI2* locus but not the *STE4* locus. We sequenced the *STE4* locus in all fifteen linked populations that had fixed a *whi2*-Q29* allele. In only two populations did the *ste4*-E315* remain heterozygous. In the other thirteen populations both the partially dominant *whi2*-Q29* mutation and the overdominant *ste4Q315\** mutation remained homozygous (**Figure S6**).

### LOH of *ste4Q315\** cannot be explained by compensatory mutations

Two possibilities could explain the observed LOH of *ste4*-E315*: either mutations at other loci on the right arm of Chromosome XV changed the net fitness effect of LOH or mutations elsewhere in the genome altered the fitness effect or the degree of dominance of either the *ste4*-E315* or the *whi2*-Q29* mutation. To test these possibilities we performed whole genome sequencing on two clones each from 20 populations: seven that lost heterozygosity at both loci (*whi2*-Q29*/*whi2*-Q29*, *ste4*-E315*/*ste4*-E315*), two that lost heterozygosity at *WHI2* but not *STE4* in *ste4*-E315* linked populations (*whi2*-Q29*/*whi2*-Q29*, *STE4*/*ste4*-E315*), nine that lost heterozygosity in a *STE4* wild-type background (*whi2*-Q29*/*whi2*-Q29*), and two control populations that are wild-type at both loci (**Figure S6**). We identified 914 nuclear mutations normally distributed across the 40 clones (**Supplemental Dataset 1**, Shapiro-Wilk test, *p=*0.36) with a mean of 30.7 mutations per clone. Twelve mutations were homozygous (not including *WHI2* or *STE4* alleles).

To identify putative *de novo* driver mutations on Chromosome XV that could explain the repeated occurrence of what should be a deleterious LOH event, we looked for homozygous nonsynonymous mutations on the right arm of Chromosome XV that are found in both clones (and thus were likely present before the LOH event). Of the 914 mutations, none meet these criteria, thus ruling out *de novo* evolution of linked beneficial mutations as an explanation for the repeated LOH of a the overdominant *ste4*-E315* mutation.

Next we looked for possible epistatic modifiers of *STE4*. Ninety-six genes either share Gene Ontology terms with *STE4* or are known physical and/or genetic interactors with *STE4* (**Table S1**). One of the of seven populations that lost heterozygosity of *ste4*-E315* acquired heterozygous missense mutations in two of these genes, *STE7* and *PTC2*. None of the other sequenced populations contained mutations in any of these 96 genes. We also took an unbiased approach by searching for genes that were mutated in more than one of the populations that lost heterozygosity of *ste4*-E315*. However, of the 101 genes containing fixed (present in both sequenced clones) nonsynonymous mutations across the 9 populations carrying an overdominant *STE4* allele, none were mutated in more than one population. In contrast, 5 of 93 genes accruing nonsynonymous fixed mutations in unlinked populations are mutated in multiple populations. Unlinked populations are significantly enriched for multi-hit genes in this comparison (Fisher’s exact, *p=*0.02).

New point mutations cannot account for LOH in all populations. To explore other possible mechanisms of modifying or escaping the overdominance of *ste4*-E315* we looked for evidence of structural evolution in our sequenced populations. First, we verified that Chromosome XV read depth in all clones is consistent with genome-wide coverage (**Figure 3A**), indicating that LOH events were due to mitotic recombination and not chromosome loss. We next identified aneuploidies and copy number variants (CNVs) in each clone. We find three different chromosomal aneuploidies across 6 populations, 5 of which were initiated with a *ste4*-E315* mutation and one of which was a control population (**Figure 3A**). Among the *ste4*-E315* populations, trisomy-VIII and trisomy-X were found in individual clones of 2 populations that lost heterozygosity at both loci. While Chromosome III aneuploidies were found in three *ste4*-E315* populations, only two of these populations lost heterozygosity at both *whi2*-Q29* and *ste4*-E315* and one lost heterozygosity at *whi2*-Q29* but retained heterozygosity at *ste4*-E315* (**Figure 3A**). We also identified two large CNVs on Chromosome III (**Table S2, Figure S7**). An amplification of 93kb on the right arm of Chromosome III, is found in both clones of a *ste4*-E315*-containing population in conjunction with a Chromosome III trisomy (**Figure 3A**). The other CNV, a 16kb deletion on the right arm of Chromosome III, was detected in a wild-type *STE4* population that lost heterozygosity at *WHI2*-Q29*.

**Figure 3.**
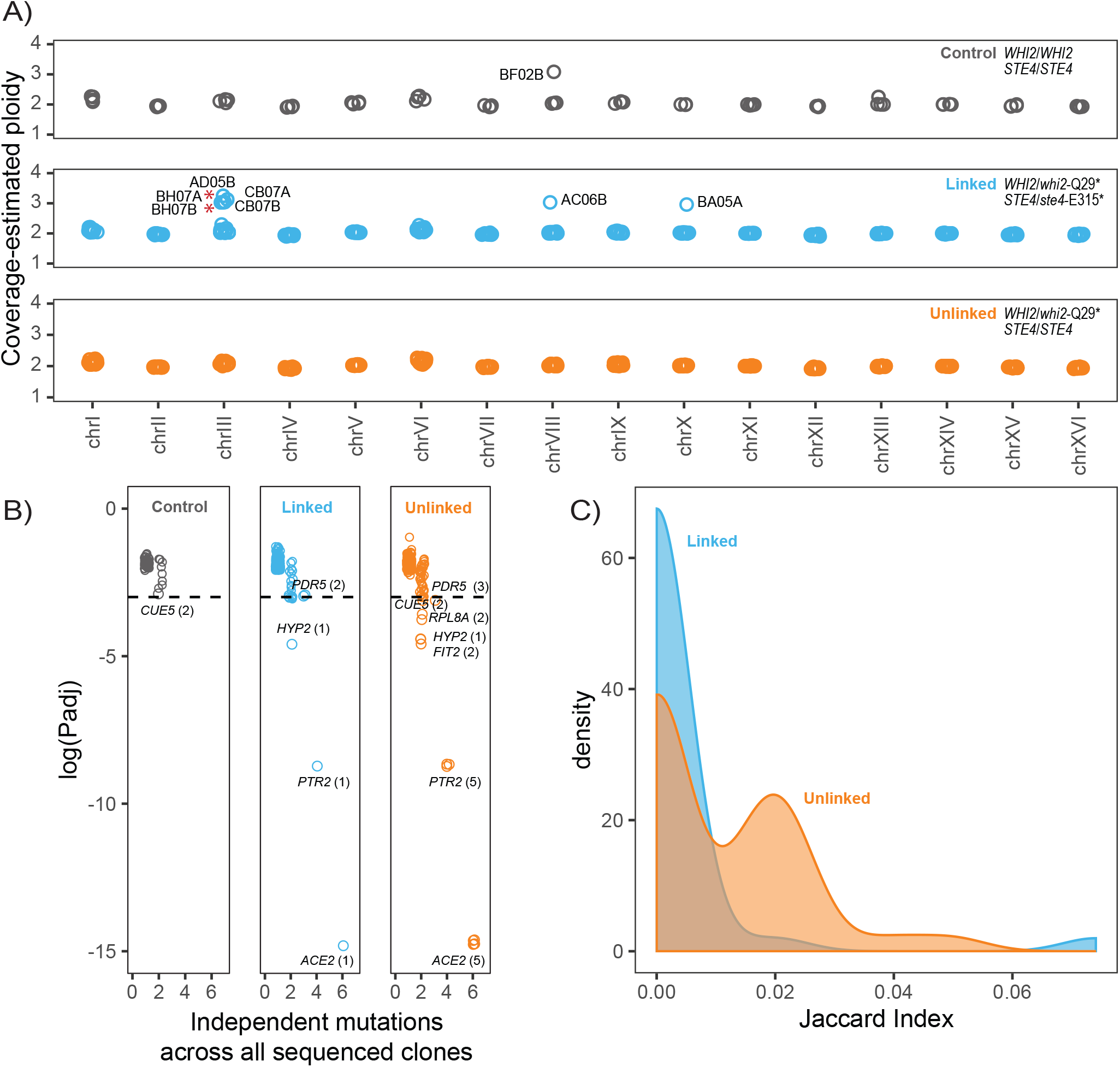
Populations with linked partially dominant and overdominant mutations show less parallelism and more varied evolutionary outcomes. **A)** Copy number of each chromosome based on median read depth in control populations (top, black), *whi2*-Q29* *ste4*-E315* populations (middle, blue), and *whi2*-Q29* *STE4* populations (bottom, orange). Chromosome XV is diploid in all populations indicating that LOH is not due to chromosome loss. Trisomy of three different chromosomes were detected. Aneuploidy was observed in 6 total populations, 5 of which contained an overdominant *STE4* mutation. Red asterisks indicate populations with Chromosome III trisomy that did not lose *ste4*-E315* heterozygosity. No aneuploidies were detected in genotypes with only *whi2*-Q29* mutations. **B)** A probability of recurrence method (6) was used to identify genes receiving significantly more mutations than expected by chance. Points are jittered on both axes to show overlapping data. A threshold of *p*<0.05 identifies seven genes as being targets of adaptive mutations in the 20 populations examined. Only 5 of the 21 total mutations in these genes occur in *ste4*-E315* linked populations. **C)** Distribution of Jaccard Indices amongst pairwise comparisons of *whi2*-Q29* *ste4*-E315* and *whi2*-Q29* *STE4* populations. The *J* distribution of linked populations is shifted significantly lower than the *J* distribution of unlinked populations (Wilcoxon rank-sum, *W*_*linked*_ =402, *W*_*unlinked*_=894, *p*<0.001). **A-C)** Points and lines are colored by genotype: (black: control, blue: *whi2*-Q29* *ste4*-E315*, orange: *whi2*-Q29* *STE4*).

There are no clear structural events shared by all populations that experienced LOH at an overdominant *STE4* allele. Chromosome III is an apparent hotspot of structural evolution in this experiment – however, most populations that underwent LOH at an overdominant locus did not contain any Chromosome III copy number variations and one of the populations that did experience Chromosome III trisomy did not lose heterozygosity at *ste4*-E315*. Consequently, we cannot attribute LOH to Chromosome III copy number variation. Nonetheless, populations carrying both *whi2*-Q29* and *ste4*-E315* alleles were enriched for aneuploidies (Fisher’s Exact, *p*=0.03) relative to populations with only a *whi2*-Q29* allele, which were all found to be euploid and had no detectable CNVs.

### Populations carrying linked dominant and overdominant beneficial mutations show a broad range of evolutionary outcomes

Taken together, we find that *ste4*-E315* *whi2*-Q29* populations are significantly depleted for recurrently mutated genes and significantly enriched for aneuploidies, a standard signature of parallel evolution. This implies that populations seeded with an overdominant mutation telomeric to a partially dominant beneficial mutation may experience a greater range of possible evolutionary outcomes. To explore this further, we aggregated all 487 mutations to coding sequences (excluding synonymous mutations) across all populations to identify targets of adaptive mutations using a recurrence-based statistical method (**Supplemental Dataset 2**). We found that only 5 of 21 mutations to 7 adaptive targets occurred in populations seeded with overdominant alleles. Since 9 of the 20 sequenced populations carried an overdominant *STE4* mutation, our null expectation would be that about half of driver mutations would be found in *ste4*-E315* populations. Instead, we find these populations to exhibit less parallelism at the gene level. We looked more closely at differences in parallelism in the set of 487 moderate to high effect coding sequence mutations by calculating the Jaccard Index (*J*) for all pairwise combinations of populations. The distribution of *J* is significantly left shifted in comparisons between *ste4*-E315\**-whi2*-Q29* populations relative to comparisons between *STE4-whi2*-Q29* populations (Wilcoxon rank-sum, *W*_*linked*_ =402, *W*_*unlinked*_ =894, *p*<0.001). Populations of both genotypes did not significantly differ in the total number of mutations accrued (*t*(15.734)= −1.68, *p*=0.112, **Figure S8)**. However, because populations lacking an overdominant *STE4* mutation tended to accrue slightly more mutations (49.4 ±12.8) than those carrying *ste4*-E315* (39.9±11.2), we used a multiple regression to show that starting genotype (*p*=0.008), but not number of mutations (*p=*0.211), is a significant predictor of *J* (F(2,69)=3.89, *p*=0.025). Therefore, we find genotypes with an overdominant *STE4* allele to be evolving more divergently at the sequence level than genotypes that are otherwise identical but lack an overdominant *STE4* allele.

## DISCUSSION

Most beneficial mutations that arise in asexually evolving diploid populations are heterozygous and are at least partially dominant. Here we show that the degree of dominance can constrain adaptive evolution at linked loci. Mechanistically these constraints arise due to conflicting effects that LOH has on partially-dominant and overdominant beneficial mutations. Given that LOH conversion tracts frequently extend the full length of a chromosome arm, the linked effects we observe on Chr XV will hold true for all chromosomes in asexually evolving diploid populations, particularly in the early stages of adaptation when overdominant beneficial mutations will be most frequent (Manna *et al*. 2011, Sellis *et al*. 2011). The strength of the linked effects, however, will vary depending on local rates of mitotic recombination and the length of repair tracts as well as the distribution of mutational effects on fitness—and the degree of dominance of those mutations—along a chromosome. Comprehensive analyses of the gene deletions in yeast reveal few underdominant or overdominant deletions (Agrawal and Whitlock), however, this may not be indicative of the distribution of dominance non-loss of function mutations. Indeed, in our evolution experiments we observe examples of overdominant, underdominant, and recessive beneficial mutations in *STE4*.

Most theory addressing mutational dominance and constraint focuses on the consequences of recessiveness, namely the constraints imposed by Haldane’s sieve (Charlesworth and Charlesworth 1999, Orr and Betancourt 2001) and the load imposed by recessive deleterious mutations (Charlesworth and Charlesworth 1999, Chasnov 2000). Underdominance is most frequently invoked as a cause of reproductive isolation (Barton and De Cara 2009), but our findings suggest an underappreciated role as an evolutionary constraint. For example, though a homozygous deletion of *STE4* would be beneficial in a *MAT***a**/**a** strain, this mutation is underdominant and would be able to fix only in extremely small or fragmented populations (Newberry *et al*. 2016).

Two of the evolved *STE4* mutations we identified demonstrate a strong degree of overdominance when engineered into an ancestral background. Recent theoretical examination of adaptation in diploids has renewed interest in the significance of overdominant mutations in adaptation and suggested overdominant polymorphisms may be a frequent mode of adaptation (Manna *et al*. 2011, Sellis *et al*. 2011). These models find that when selection on a trait is stabilizing, strong effect heterozygous mutations that overshoot the fitness optimum as homozygotes should be somewhat common. Experimental evolution provides a way to empirically test this prediction. The few examples of overdominance arising *de novo* in laboratory evolution include amplifications of glucose transporter genes in glucose-limited chemostat populations (Sellis *et al*. 2016). Overdominance of a copy number variant is well explained by an “overshoot” of an optimal gene copy number, and hexose transporter amplifications have been previously shown to exhibit sign epistasis with mutations that upregulate their expression.

Mitotic recombination resulting in loss-of-heterozygosity (LOH) is a common and important mechanism of adaptation in laboratory evolving diploid yeast (Gerstein *et al*. 2014, Smukowski Heil *et al*. 2017, Fisher *et al*. 2018, James *et al*. 2019). Most of these reported instances of LOH in asexual yeast adaptation involve a large conversion tract that runs from the break point to the telomere. This means that there is effective linkage between loci that are kilobases apart. Overdominant beneficial mutations will therefore impose constraint on mitotic recombination along the full length of a chromosome arm. We tested this using a partially dominant beneficial mutation 700 kb upstream of *STE4* in the *WHI2* gene to examine how LOH dynamics differ between genotypes with only a *WHI2* mutation and those with a *WHI2* mutation linked to an overdominant *STE4* allele. In order to lose heterozygosity and gain a fitness benefit at *WHI2*, linked populations that must either lose heterozygosity at the *WHI2* locus while maintaining heterozygosity at the distal *STE4* locus or suffer the fitness cost loss of gene-converting an overdominant *STE4* locus. This is the first evolution experiment to directly measure rates of LOH when the conversion of a linked locus is unfavorable. We find a significant initial obstructive effect of overdominant mutations on the rate of adaptive LOH at linked loci. After the first few hundred generations, however, this effect is weakened and LOH is repeatedly observed in populations bearing an overdominant *STE4* allele.

Given the individual fitness effects of *whi2*-Q28* and *ste4*-Q315* alleles, we expected fewer populations to lose heterozygosity on Chromosome XV when both mutations were present on the same chromosome. We found, however, that linked populations still adapted by way of LOH at *WHI2*, but the appearance of these events was delayed by several hundred generations (because we were only able to detect LOH when homozygous genotypes were above 50% and could not observe initial appearance of these homozygotes in the populations). One possible explanation is that *de novo* mutations arose during this time that changed the net fitness effect of LOH. Whole genome sequencing revealed that, while modifying mutations may occur, they cannot be the sole explanation for the LOH we observe in the linked populations. We do observe an enrichment for aneuploidies in the linked populations, though the specific changes we found (gains of additional copies of Chromosomes III and VIII) have been observed before (Fisher *et al*. 2018).

Analysis of genome sequence evolution in 20 sequenced clones shows that unlinked *whi2*-Q28* populations accrue more mutations in common targets of selection, whereas *whi2*-Q28*/*ste4*-Q315* linked populations show a wider range of evolutionary outcomes at the genome sequence level. The difference in the modes of adaptation between two nearly identical genomes (differing only by a single heterozygous mutation), indicate that small changes in the genome can introduce constraints on genome evolution and influence evolutionary outcomes.

## METHODS

### Analysis of evolved mutations in *STE4*

We previously identified *STE4* mutations from whole-genome sequencing data reported for 40 haploid (Lang *et al*. 2013) and 46 autodiploid (Fisher *et al*. 2018) yeast populations. The mutational target window of *STE4* (in bp) was calculated for both haploids and autodiploids. The probability of all mutations occurring in the observed window was calculated separately for haploids and autodiploids using a one-sample proportions test.

Homology modeling of Ste4p was performed automatically on the SWISS-MODEL web server to visualize the positions of mutated residues. The best scoring model was based on the structure of G protein subunit beta (Gnb1) from *Rattus norvegicus* (41.96% identity, 0.63 GMQE, −3.31 QMEAN, PDB ID: 6CMO). Visualization of the Ste4p model was done in PyMOL Molecular Graphics System, version 2.3.0. This structure does not contain a 33 amino acid yeast-specific insertion.

### Construction of evolved mutation and *STE4* deletion strains

Strains used in this paper are described in **Table S3**. Reconstruction experiments were performed in the same W303 ancestral background (yGIL432; *MAT***a**, *ura3*Δ::p*FUS1*-yEVenus, *ade2-1, his3-11,15, leu2-3,112, trp1-1, CAN1, bar1*Δ::*ADE2, hmlα*Δ::*LEU2, GPA1*::NatMX). Briefly, deletion strains were generated by integrating the *ste4*Δ::KanMX locus from the deletion collection. Crosses of strains carrying null alleles were performed by first transforming with a *STE4*-expressing plasmid from the MoBY ORF plasmid collection to compliment *ste4*Δ. Three evolved *STE4* alleles were selected for reconstruction, 81ΔT (S261fs), G943T (E315*), and G935A (R312Q). Alleles were reconstructed in yGIL432 using CRISPR-Cas9 alleles swaps. We constructed constitutive Cas9-expressing plasmids starting from pML104 (Addgene 67638) expressing a *STE4*-specific guide RNAs (5’ CTACCCCTAC TTATATGGCA 3’) and co-transformed the plasmid along with a 500 bp linear repair template (gBlock, IDT) encoding the one of three evolved alleles as well as a synonymous C954A PAM site substitution. A strain containing just the synonymous PAM site was also isolated to verify neutrality (**Figure S3**). To minimize variation due to transformation and Cas9 activity, one successful transformant per allele was backcrossed twice and the resulting diploid was sporulated and tetrad dissected. For each allele, spores were genotyped at *STE4* and intercrossed to generate heterozygous and homozygous mutants. Crosses of strains carrying evolved *ste4* alleles were performed by first transforming with a plasmid from the MoBY ORF plasmid collection to compliment *STE4*. Mutants carrying an evolved *whi2*-C85T (Q29*) allele were generated in identical fashion with two exceptions. The evolved substitution is within the *WHI2* gRNA used (5’ ACAGTACGAA GGTAACGAGG 3’), and therefore no synonymous mutation was introduced to eliminate Cas9 activity. A correct *whi2*-Q29* was backcrossed once and intercrossed to produce homozygotes and heterozygotes. All diploid genotypes were converted to *MAT***a**/**a** as described above. We also generated strains containing dominant drug cassettes tightly linked to the *WHI2* locus using CRISPR. We inserted either HphMX or KanMX 220 bp downstream of *WHI2* or *whi2*-Q29* by transforming with the same gRNA (5’ ATCCCCTTCT GCAAATAACG 3’) and Cas9-expressing plasmid and co-transforming with linear drug cassettes flanked by 40 bp of homology to the targeted region. Successful transformants were then backcrossed to either a wild-type background or a *ste4*-G943T (described above) mutant. Crosses were sporulated and spores were selected in which drug-marker tagged mutant and wild-type *WHI2* alleles are present on the same chromosome as both mutant and wild-type *STE4*. Correct spores were crossed to generate three genotypes: *WHI2*::HphMX/*WHI2*::KanMx *STE4*/*STE4, WHI2*::HphMX/*whi2*-Q29*::KanMx *STE4*/*ste4*-E315*, and *WHI2*::HphMX/*whi2*-Q29*::KanMx *ste4*-E315*/*STE4*. All three genotypes were converted to *MAT***a**/**a** as described above. Eight replicate *MAT***a**/**a** colonies were picked for each mating-type conversion to be used for downstream analysis.

### Fitness assays

We measured the effects of complete gene deletions and evolved *STE4* mutations on fitness using competitive fitness assays as previously reported (Buskirk *et al*. 2017). Briefly, query cultures were mixed 1:1 with a ploidy and mating-type matched fluorescently labeled ancestral strain. Co-cultures were propagated in a 96-well plate in an identical to the evolution experiment in in which the variants arose for 50 generations. Saturated cultures were sampled for flow cytometry at ten-generation intervals. Flow cytometry data was analyzed with FlowJo 10.3. Selective coefficients were calculated as the slope of the best-fit line of the natural log of the ratio between query and reference strains against time.

Two technical replicates each of eight biological replicates were averaged for analysis of all *MAT***a**/**a** genotypes and all deletion mutants. Evolved mutations in a haploid background were averaged from four technical replicates of a single clone. Fitness data for haploid and diploid genotypes were analyzed independently using a one-way analysis of variance ANOVA. Post hoc comparisons using the Tukey test were carried out to identify genotypes with significantly different fitness than wild-type controls.

### Short-term evolution experiment

We examined the effect of evolved *ste4* alleles on likelihood of loss-of-heterozygosity (LOH) at a linked locus, *WHI2*. We first validated the homozygous and heterozygous fitness benefits of an evolved allele, *whi2*-C85T (Q29*), via mutant reconstruction and fitness assays as described above. We then generated strains containing dominant drug cassettes tightly linked to the *WHI2* locus to investigate the effect of *STE4* linkage on loss of heterozygosity along the right arm of Chromosome XV (Supplementary methods).

Three strains (*WHI2*::HphMX/*WHI2*::KanMx *STE4*/*STE4, WHI2*::HphMX/*whi2*-Q29*::KanMx *STE4*/*ste4*-E315*, and *WHI2*::HphMX/*whi2*-Q29*::KanMx *ste4*-E315*/*STE4*) were grown in 10 ml overnight cultures in YPD with 0.4 mg/ml G418 and 0.6 mg/ml Hygromycin B. Saturated cultures were diluted 1:1,000 to initiate 287 128 μl cultures across three 96-well plates. Plates were incubated unshaken at 30°C and propagated daily in an identical fashion to the original evolution experiment in which the mutations arose (Lang *et al*. 2013). After 500 generations heterozygosity was assayed by spotting 2 μl (∼5,000 cells) to double drug plates (YPD with 0.4 mg/ml G418, 0.6 mg/ml Hygromycin B) and to both single drug agar plates. Plates were inspected for speckled spots (indicating homozygous genotypes in ≥50% the population) and absence of growth (indicating LOH sweeps). We compared the number of populations with evidence of LOH polymorphism or sweeps between genotypes using a Fisher’s exact test with a Bonferroni correction for multiple comparisons.

### Whole genome sequencing and analysis

We sequenced nine of fifteen linked populations and nine of twenty-seven unlinked populations in which a homozygous *WHI2* allele genotype fixed. The nine populations for each group were chosen to be representative of the spectrum of dynamics observed in the evolution experiment (i.e. some populations that underwent LOH early in the experiment and some that underwent LOH late). Each population was struck to singles on YPD to obtain two clones for sequencing. Clones were grown overnight in 5 ml YPD and then frozen as cell pellets at −20°C. Genomic DNA was isolated from frozen pellets via phenol-chloroform extraction and ethanol precipitation. Total genomic DNA was used in a Nextera library preparation as described previously (Buskirk *et al*. 2017). All individually barcoded clones were pooled and paired-end sequenced on NovaSeq 6000 sequencer at the Genomics Core Facility at the Lewis-Sigler Institute for Integrative Genomics, Princeton University.

Raw sequence data were concatenated and then demultiplexed using a custom python script from L. Parsons (Princeton University). Adapter sequences were removed using Trimmomatic (Bolger *et al*. 2014). Reads were then aligned to a customized W303 genome using BWA v. 0.7.7 (Li and Durbin 2009) and variants were called using FreeBayes v0.9.21-24-381. VCFtools was used to filter variants common to all genomes. Remaining mutations were annotated using a strain-background customized annotation file (Matheson *et al*. 2017). All putative evolved mutations were confirmed manually using IGV (Thorvaldsdóttir *et al*. 2013).

Each genome was independently examined for structural variants using Samtools depth (Li *et al*. 2009). Aneuploidies were detected by dividing median chromosome coverage by median genome-wide coverage for each chromosome. CNVs were similarly detected using a sliding 1 kb window across each chromosome. Putative CNVs identified were confirmed by visual inspection of chromosome coverage plots.

A list of 92 candidate genes for the modification of overdominance at *STE4* was curated by concatenating a list of all known *STE4* interactors and all genes annotated to the GO term “pheromone dependent signal transduction involved in conjugation with cellular fusion” and all genes annotated to children of this GO term. The above GO term was selected out of the seventeen terms assigned to *STE4* because changes to pheromone-induced signaling is thought to be the cellular basis of the fitness effect of *STE4* mutations (Lang *et al*. 2009). These searches were performed in YeastMine (Balakrishnan *et al*. 2012).

### Identification of genic targets of selection and quantification of parallelism

To identify parallel targets of selection we first removed all non-protein coding and synonymous mutations to improve our signal. We then identified parallel targets of selection as described previously (Fisher *et al*. 2018). Briefly, we calculate the expected number of mutations for each gene, σ, using the Poisson distribution weighted by the length (*L*) of the gene in base-pairs:

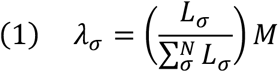

where *M* is the total number of coding sequence mutations in the dataset. The probability of observing *k* mutations in gene *σ* is therefore

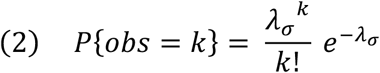

We use expression (2) to calculate the *p*-value for the observed number of mutations in each gene. We then applied a Benjamin-Hochberg post hoc adjustment to correct for multiple hypothesis testing.

Parallelism was quantified using the Jaccard Index (Bailey *et al*. 2015), which calculates the similarity between two sets of mutated genes by quantifying the overlap of their union. Again, non-protein coding and synonymous mutations were excluded to increase the signal of adaptive parallelism. A value of *J* was calculated for pairwise combinations of populations. The distribution of *J* for all pairs of linked populations was compared to the distribution of *J* for all pairs of unlinked populations using a Wilcoxon rank-sum test with continuity correction.

### Statistical analyses

All statistical analyses reported were performed using tools in the R Stats package in R v.3.6.2. All plots were produced in R using the ggplot2 package (Wickham 2016) except Figures S5 and S7 which were produced using base R plotting.

## Supporting information

Combined Supplemental Figures 1-8

## ACKNOWLEDGEMENTS

We thank members of the Lang Lab for comments on this manuscript. This work was supported by the National Institutes of Health Grant 1R01GM127420 [G.I.L.]. K.J.F. is a Morgridge Metabolism Interdisciplinary Fellow of the Morgridge Institute for Research.

## AUTHOR CONTRIBUTIONS

K.J.F. and G.I.L. conceived of the project and designed experiments. K.J.F. and R.C.V. performed experiments. K.J.F. and R.C.V. analyzed the data. K.J.F. and G.I.L. wrote the manuscript.

## DATA AVAILABILITY

The raw short-read sequencing data reported in this paper have been deposited in the NCBI BioProject database (accession no. PRJNA634573).

