## Supplementary material for "Overdominant mutations restrict adaptive loss of heterozygosity at linked loci": Combined Supplemental Figures 1-8

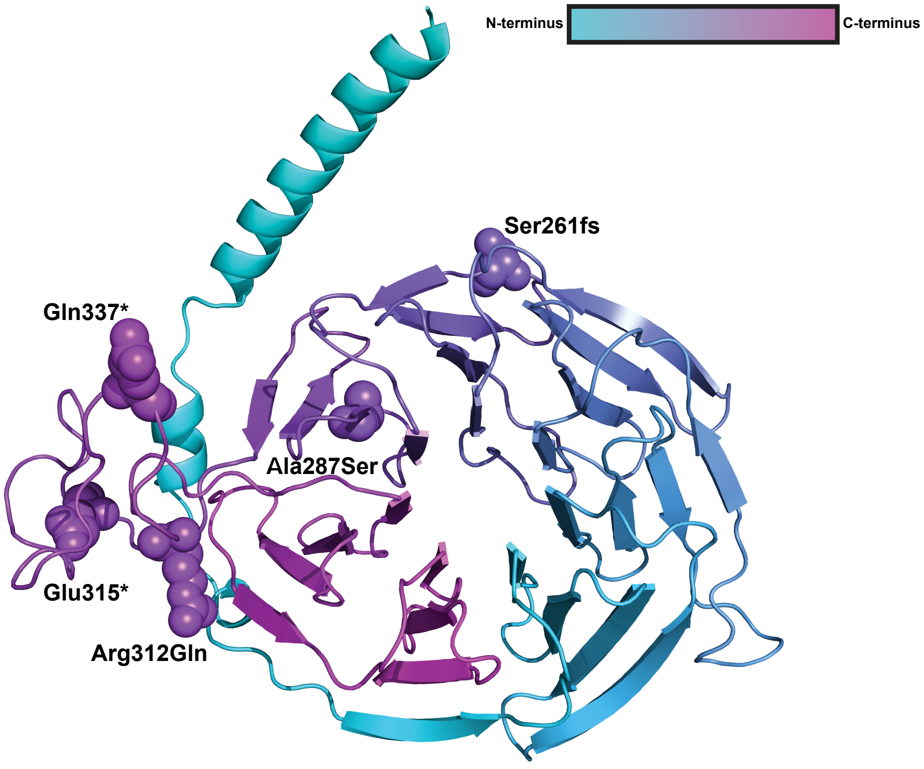


**Figure S1.** Homology-based model of Ste4. Six *STE4* mutations (Gly250Gly, Ser261fs, Ala287Ser, Arg312Gln, Glu315*, Gln337*) arose among 46 autodiploid populations (Fisher *et al.* 2018). The five non-synonymous mutations are labeled and shown as spheres. R312Q, E315*, and E337* are contained within the yeast-specific putative random coil region.

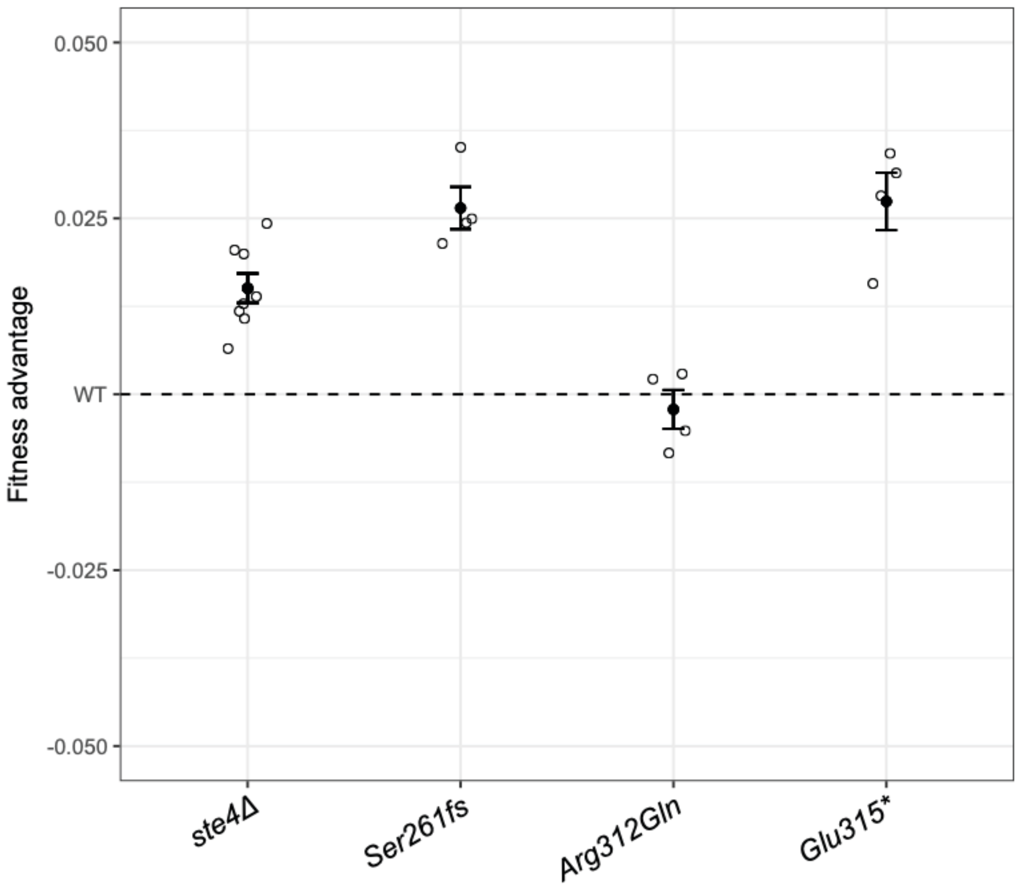


**Figure S2.** Fitness effects of *ste4*Δ and evolved mutations in a haploid background. The data confirm a previously reported benefit of the knockout (Lang *et al.* 2009). Two of the three autodiploid-evolved mutations have a significant fitness benefit in a haploid background while one, the only non-truncation allele, appears neutral. Open points represent selection coefficients from eight biological replicates for *ste4*Δ and four biological replicates of each evolved genotype. Bold point is the mean ± standard error.

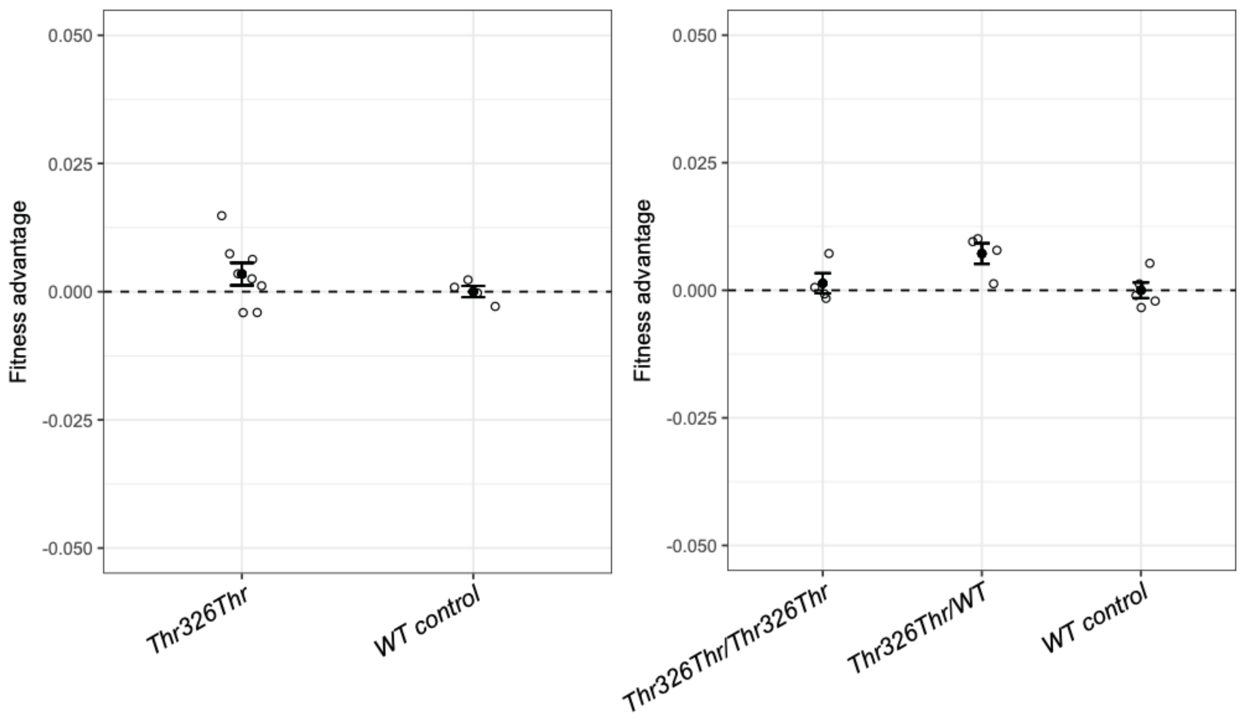


**Figure S3.** A synonymous Thr326Thr mutation was introduced along with each evolved mutation during Cas9-mediated allele swaps to ablate the PAM site of the gRNA target sequence. Single mutants for the synonymous mutation were isolated and assayed along with double mutants carrying evolved mutations. Synonymous mutant fitness did not differ from wild type in a haploid background ($p$=0.93), a heterozygous diploid background ($p$=0.24), or a homozygous diploid background ($p$=1.0). Open points represent selection coefficients from eight biological replicates for the mutation in a haploid background and four biological replicates of each diploid genotype. Bold point is the mean ± standard error.

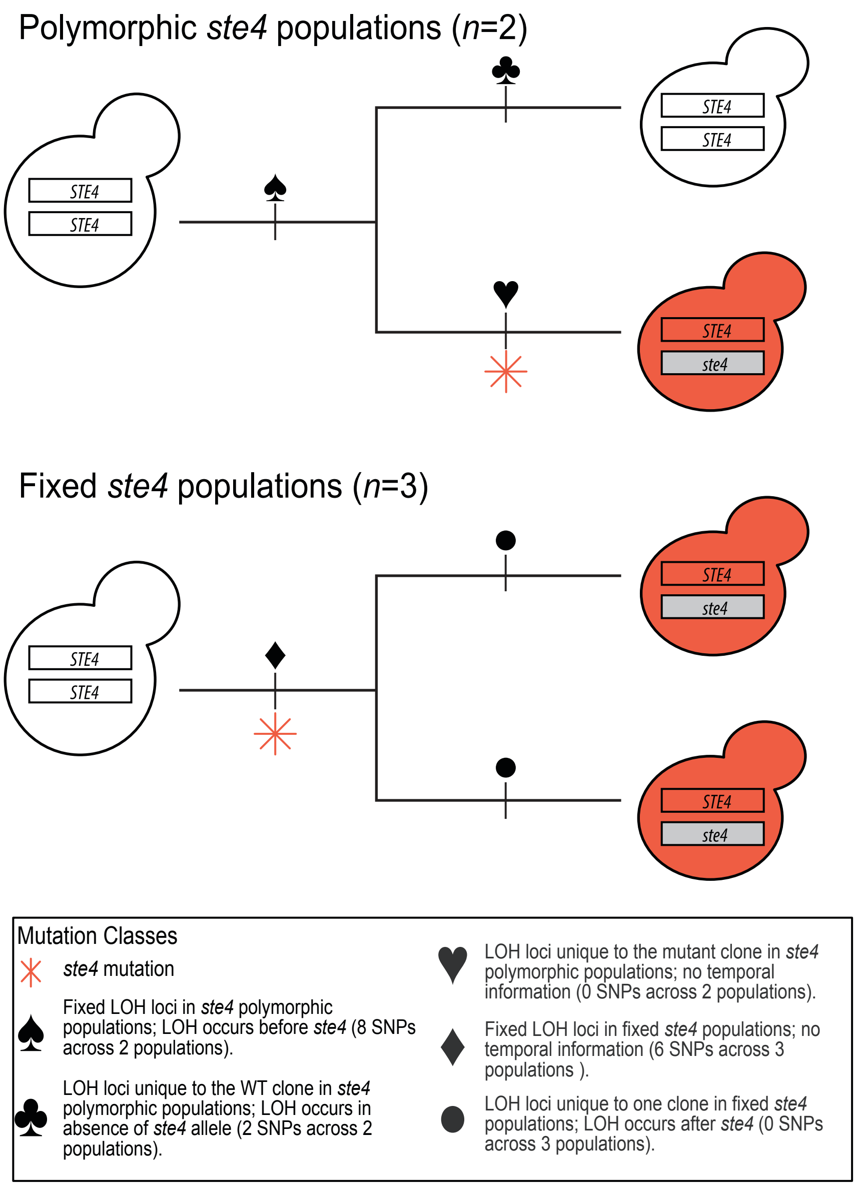


**Figure S4**. Standard curve for LOH detection. Heterozygosity was tracked by automated spotting of a 2 µl volume containing ~5,000 cells per population to double drug (0.4 mg/ml G418, 0.6 mg/ml Hygromycin B) YPD agar one-well plates. Spotting a series of known populations to double drug media shows that speckling becomes evident once homozygous genotypes reach 50% and above. Populations were also spotted in parallel to both single drugs to determine which *WHI2* allele had lost heterozygosity.

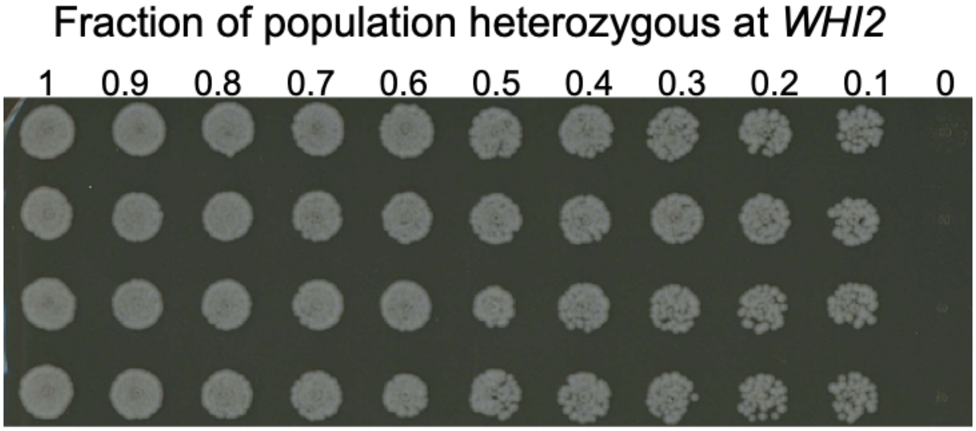


**Figure S5.** Time-course dynamics of heterozygosity at the *WHI2* locus in all 57 populations that showed evidence of LOH. Lines are colored by treatment group (green: linked, yellow: unlinked, blue: control). The portion of the population homozygous at *WHI2* was estimated by spotting populations to double and single drug media. Some populations (A_C09, A_D09, A_G10, A_G11, A_H09, A_H11, B_C11, B_F09, C_A12, C_B11, C_D11, and C_F10) had already experienced LOH at the first time point surveyed (Generation 420). Linked populations with bolded lines indicate populations with fixed homozygous *whi2*-Q29*. Dashed lines indicate the two linked that populations that retained heterozygosity at *STE4.* Populations from which 2 clones were sequenced are indicated by asterisks.

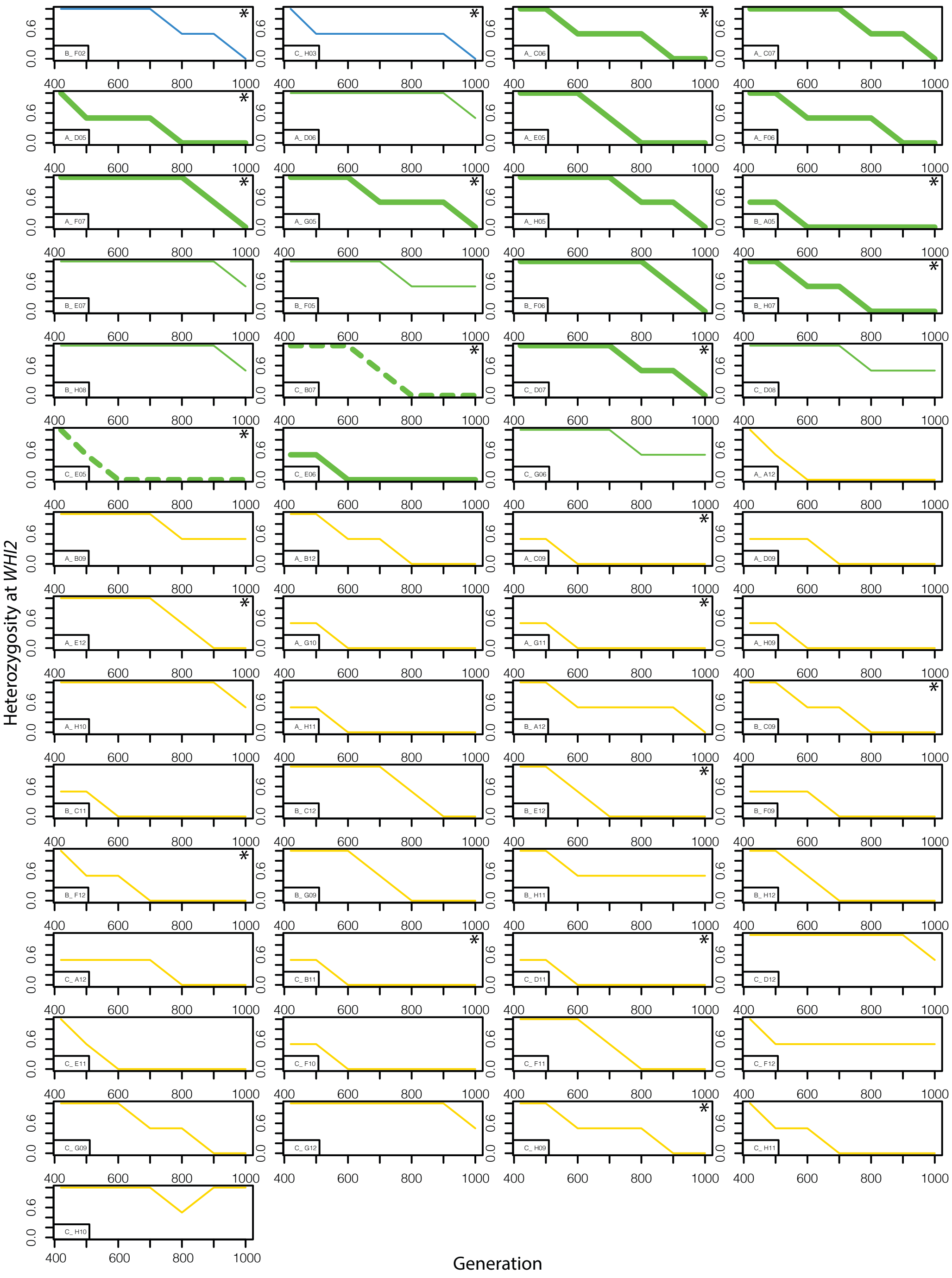


**Figure S6.** We examined endpoint sequencing data from the experiment in which overdominant *STE4* mutations evolved to look for any evidence of LOH occurring on the right arm of Chromosome XV after a *STE4* mutation established. *STE4* mutations arose in 6 independent populations. In 2 of these, the mutation is unique to one of two sequence clones (polymorphic). In 3 populations the *STE4* mutation is shared by both sequenced clones (fixed). Only one clone of the remaining population was sequenced, and this population provides no temporal information and is not included in the figure. We found 8 fixed homozygous SNPs in the 2 polymorphic *ste4* populations – providing evidence that LOH occurs before *STE4* mutations arise (spades). Conversely, we found 0 homozygous loci that were unique to one clone in fixed *ste4* populations (circles) and therefore no evidence that LOH occurred in any populations after the establishment of *STE4* mutations. The other 3 classes of mutations are indicated but provide no temporal information.

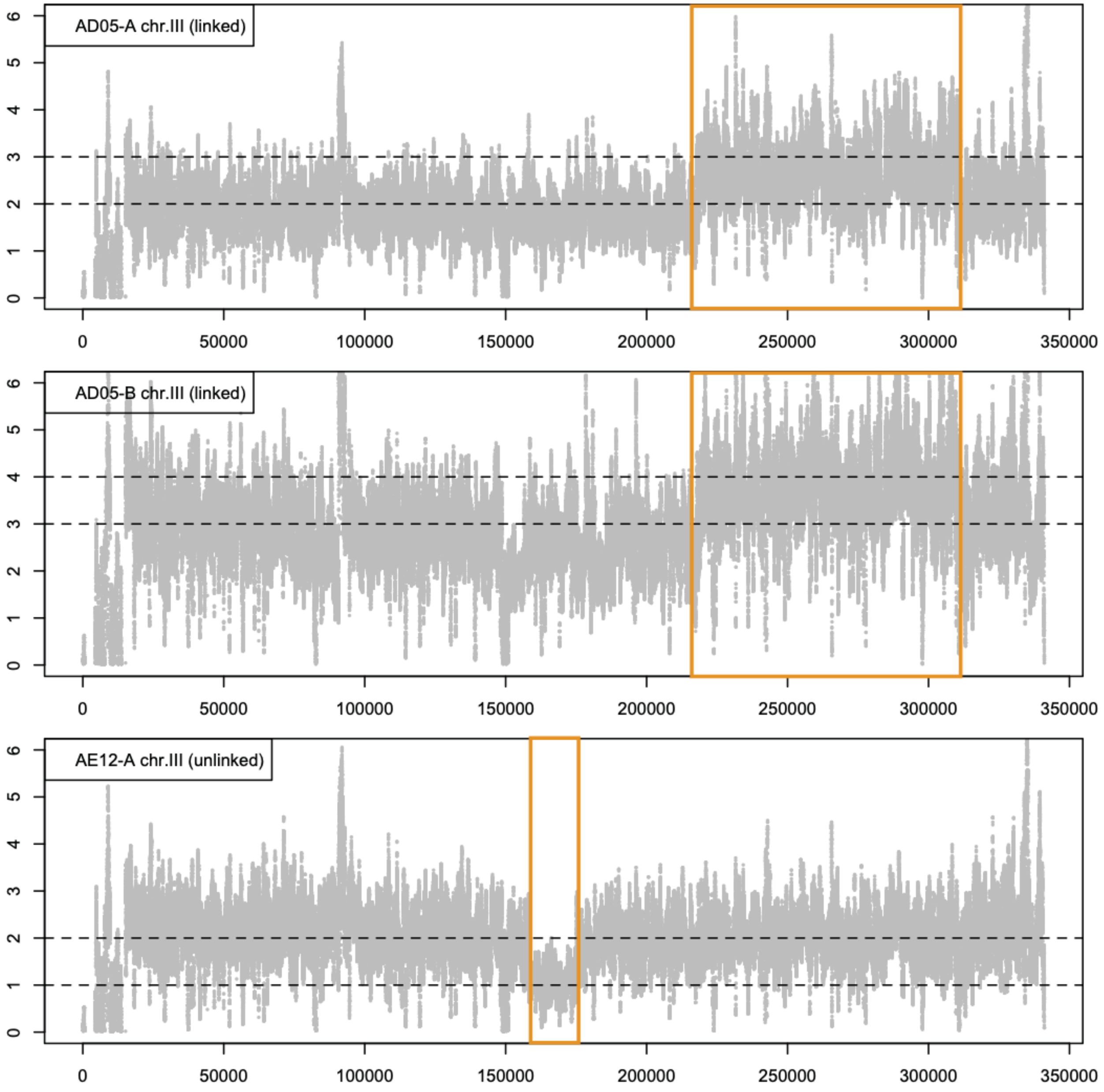


**Figure S7.** We detected copy number variants by examining sliding 1 kb windows along each chromosome. Windows with suspected elevated or decreased coverage were visually inspected using coverage plots. A 95 kb amplification in both clones from a linked population and a 16 kb deletion in one clone from an unlinked population were both identified on the right arm of Chromosome III. The dotted lines indicate the ploidy of the chromosome (ADO5-B has a Chromosome III trisomy) and the copy number of the amplified or deleted region. Orange boxes indicate the windows in which the CNVs were found.

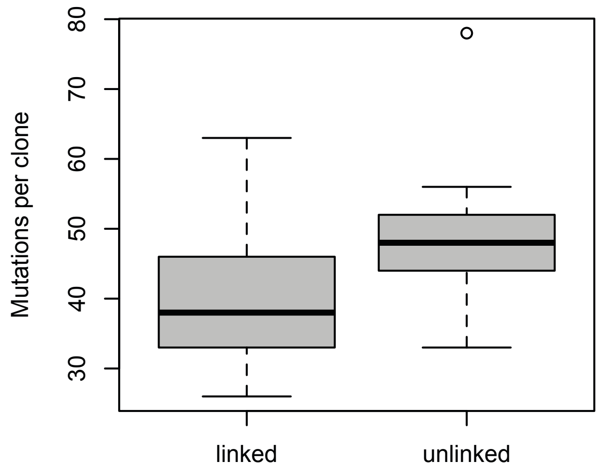


**Figure S8.** Boxplot showing the distributions of mutations per population by starting genotype. There is no significant difference in accumulated mutations per population between genotypes ($t$(15.734)= -1.68, $p$=0.11).

**Table S1 Known physical and/or genetic interactors, and genes sharing Gene Ontology terms with STE4**

| Gene | Systematic Name | Populations | Gene | Systematic Name | Populations |
|---|---|---|---|---|---|
| CLN3 | YAL040C | NA | SAP185 | YJL098W | NA |
| CDC24 | YAL041W | NA | FAR1 | YJL157C | NA |
| KIN3 | YAR018C | NA | CCT5 | YJR064W | NA |
| FUS3 | YBL016W | NA | MOG1 | YJR074W | NA |
| PKC1 | YBL105C | NA | STE18 | YJR086W | NA |
| TEC1 | YBR083W | NA | PMT4 | YJR143C | NA |
| BEM1 | YBR200W | NA | ELM1 | YKL048C | NA |
| RNQ1 | YCL028W | NA | STE3 | YKL178C | NA |
| STE50 | YCL032W | NA | CNB1 | YKL190W | NA |
| CDC39 | YCR093W | NA | PUF3 | YLL013C | NA |
| ARP2 | YDL029W | NA | HOG1 | YLR113W | NA |
| STE7 | YDL159W | AF07 | STE11 | YLR362W | NA |
| DHH1 | YDL160C | NA | SST2 | YLR452C | NA |
| CDC36 | YDL165W | NA | MCM1 | YMR043W | NA |
| PTP1 | YDL230W | NA | STV1 | YMR054W | NA |
| STE5 | YDR103W | NA | PGM2 | YMR105C | NA |
| PLP1 | YDR183W | NA | ADE17 | YMR120C | NA |
| CCT6 | YDR188W | NA | DSK2 | YMR276W | NA |
| TCP1 | YDR212W | NA | FCP1 | YMR277W | NA |
| AHA1 | YDR214W | NA | MSG5 | YNL053W | NA |
| AKR1 | YDR264C | NA | OCA2 | YNL056W | NA |
| SSF2 | YDR312W | NA | YAF9 | YNL107W | NA |
| MSN5 | YDR335W | NA | SKO1 | YNL167C | NA |
| RAD23 | YEL037C | NA | MDG1 | YNL173C | NA |
| GCD11 | YER025W | NA | SSB2 | YNL209W | NA |
| MOT2 | YER068W | NA | RPD3 | YNL330C | NA |
| PTC2 | YER089C | AF07 | NUF2 | YOL069W | NA |
| DSE1 | YER124C | NA | RGA1 | YOR127W | NA |
| RSP5 | YER125W | NA | PTP2 | YOR208W | NA |
| STE2 | YFL026W | NA | PLP2 | YOR281C | NA |
| SEC53 | YFL045C | NA | PPQ1 | YPL179W | NA |
| NAB2 | YGL122C | NA | SSO1 | YPL232W | NA |
| KSS1 | YGR040W | NA | CLN2 | YPL256C | NA |
| ESP1 | YGR098C | NA | LTP1 | YPR073C | NA |
| RSR1 | YGR152C | NA | RHO1 | YPR165W | NA |
| STE20 | YHL007C | NA | DPM1 | YPR183W | NA |
| SBP1 | YHL034C | NA | STE4 | YOR212W | NA |
| GPA1 | YHR005C | NA | SCP160 | YJL080C | NA |
| YHR033W | YHR033W | NA | PTC1 | YDL006W | NA |
| SSF1 | YHR066W | NA | MFA2 | YNL145W | NA |
| STE12 | YHR084W | NA | MFA1 | YDR461W | NA |
| YCK1 | YHR135C | NA | MF(ALPHA)2 | YGL089C | NA |
| BAR1 | YIL015W | NA | MF(ALPHA)1 | YPL187W | NA |
| SYG1 | YIL047C | NA | GET3 | YDL100C | NA |
| POG1 | YIL122W | NA | FYV5 | YCL058C | NA |
| CCT2 | YIL142W | NA | CDC42 | YLR229C | NA |
| CCT3 | YJL014W | NA | ADF1 | YCL058W-A | NA |
| MTR4 | YJL050W | NA | | | |

**Table S2. Chromosome III CNVs**

| Chromosome | Start | Stop | Size(kb) | Copy Number | Type | Population | Group | Genes contained in CNV |
|---|---|---|---|---|---|---|---|---|
| chrIII | 159000 | 175000 | 16 | 1 | loss | AE12 A | unlinked | MAK32, PET18, MAK31, HTL1, HSP30, YCR023C, SLM5, PMP1, NPP1, RHB1 |
| chrIII | 218000 | 311000 | 93 | 3 | gain | AD05 A | linked | HMRA1, MATALPHA1, TAF2, YCR043C, PER1, YCR045C, IMG1, BUD23, ARE1, YCR051W, RSC6, THR4, CTR86, PWP2, YIH1, TAH1, YCR061W, tS(CGA)C, tS(CGA)C, BUD31, HCM1, RAD18, SED4, ATG15, CPR4, IMG2, RSA4, SSK22, SOL2,ERS1, YCR075W-A, YCR076C, PAT1, PTC6, SRB8, AHC2, TRX3, TUP1, CSM1, YCR087C-A, ABP1, FIG2, YCR090C, KIN82, MSH3, CDC39, CDC50,YCR095C |
| chrIII | 218000 | 311000 | 93 | 4* | gain | AD05 B | linked | HMRA1, MATALPHA1, TAF2, YCR043C, PER1, YCR045C, IMG1, BUD23, ARE1, YCR051W, RSC6, THR4, CTR86, PWP2, YIH1, TAH1, YCR061W, tS(CGA)C, tS(CGA)C, BUD31, HCM1, RAD18, SED4, ATG15, CPR4, IMG2, RSA4, SSK22, SOL2,ERS1, YCR075W-A, YCR076C, PAT1, PTC6, SRB8, AHC2, TRX3, TUP1, CSM1, YCR087C-A, ABP1, FIG2, YCR090C, KIN82, MSH3, CDC39, CDC50,YCR095C |

* This clone has also has ChrIII trisomy

**Table S3. Strains used in this paper**

| Strain | Ploidy | Assay | Genotype*† |
|---|---|---|---|
| yGIL1120 | MATa | Competitive fitness | STE4, ade2–1, his3–11, leu2–3,112, trp1–1, ura3Δ::PFus1-yEVenus, bar1Δ::ADE2, hmlαΔ::LEU2, GPA1::NatMX |
| yGIL1121 | MATa | Competitive fitness | ste4 Δ::KanMX, ade2–1, his3–11, leu2–3,112, trp1–1, ura3Δ::PFus1-yEVenus, bar1Δ::ADE2, hmlαΔ::LEU2, GPA1::NatMX |
| yGIL1611 | MATa/a | Competitive fitness | STE4, ade2–1, his3–11, leu2–3,112, trp1–1, ura3Δ::PFus1-yEVenus/ura3, bar1Δ::ADE2, hmlαΔ::LEU2, GPA1::NatMX |
| yGIL1612 | MATa/a | Competitive fitness | STE4/ ste4 Δ::KanMX, ade2–1, his3–11, leu2–3,112, trp1–1, ura3Δ::PFus1-yEVenus/ura3, bar1Δ::ADE2, hmlαΔ::LEU2, GPA1::NatMX |
| yGIL1613 | MATa/a | Competitive fitness | ste4 Δ::KanMX, ade2–1, his3–11, leu2–3,112, trp1–1, ura3Δ::PFus1-yEVenus/ura3, bar1Δ::ADE2, hmlαΔ::LEU2, GPA1::NatMX |
| yGIL1619 | MATa | Competitive fitness | ste4C958A‡, ade2–1, his3–11, leu2–3,112, trp1–1, ura3Δ::PFus1-yEVenus, bar1Δ::ADE2, hmlαΔ::LEU2, GPA1::NatMX |
| yGIL1642 | MATa/a | Competitive fitness | ste4C958A‡/STE4, ade2–1, his3–11, leu2–3,112, trp1–1, ura3Δ::PFus1-yEVenus/ura3, bar1Δ::ADE2, hmlαΔ::LEU2, GPA1::NatMX/ GPA1::KanMX |
| yGIL1643 | MATa/a | Competitive fitness | ste4C958A‡/ ste4C958A‡, ade2–1, his3–11, leu2–3,112, trp1–1, ura3Δ::PFus1-yEVenus/ura3, bar1Δ::ADE2, hmlαΔ::LEU2, GPA1::NatMX/ GPA1::KanMX |
| yGIL1616 | MATa | Competitive fitness | ste4T81Δ, ade2–1, his3–11, leu2–3,112, trp1–1, ura3Δ::PFus1-yEVenus, bar1Δ::ADE2, hmlαΔ::LEU2, GPA1::NatMX |
| yGIL1636 | MATa/a | Competitive fitness | ste4T81Δ/STE4, ade2–1, his3–11, leu2–3,112, trp1–1, ura3Δ::PFus1-yEVenus/ura3, bar1Δ::ADE2, hmlαΔ::LEU2, GPA1::NatMX/GPA1::KanMX |
| yGIL1637 | MATa/a | Competitive fitness | ste4T81Δ, ade2–1, his3–11, leu2–3,112, trp1–1, ura3Δ::PFus1-yEVenus/ura3, bar1Δ::ADE2, hmlαΔ::LEU2, GPA1::NatMX/GPA1::KanMX |
| yGIL1617 | MATa | Competitive fitness | ste4G943T, ade2–1, his3–11, leu2–3,112, trp1–1, ura3Δ::PFus1-yEVenus, bar1Δ::ADE2, hmlαΔ::LEU2, GPA1::NatMX |
| yGIL1638 | MATa/a | Competitive fitness | ste4G943T /STE4, ade2–1, his3–11, leu2–3,112, trp1–1, ura3Δ::PFus1-yEVenus/ura3, bar1Δ::ADE2, hmlαΔ::LEU2, GPA1::NatMX/GPA1::KanMX |
| yGIL1639 | MATa/a | Competitive fitness | ste4G943T, ade2–1, his3–11, leu2–3,112, trp1–1, ura3Δ::PFus1-yEVenus/ura3, bar1Δ::ADE2, hmlαΔ::LEU2, GPA1::NatMX/GPA1::KanMX |
| yGIL1618 | MATa | Competitive fitness | ste4G935A, ade2–1, his3–11, leu2–3,112, trp1–1, ura3Δ::PFus1-yEVenus, bar1Δ::ADE2, hmlαΔ::LEU2, GPA1::NatMX |
| yGIL1640 | MATa/a | Competitive fitness | ste4G935A /STE4, ade2–1, his3–11, leu2–3,112, trp1–1, ura3Δ::PFus1-yEVenus/ura3, bar1Δ::ADE2, hmlαΔ::LEU2, GPA1::NatMX/GPA1::KanMX |
| yGIL1641 | MATa/a | Competitive fitness | ste4G935A, ade2–1, his3–11, leu2–3,112, trp1–1, ura3Δ::PFus1-yEVenus/ura3, bar1Δ::ADE2, hmlαΔ::LEU2, GPA1::NatMX/GPA1::KanMX |
| yGIL1555 - control | MATa/a | Evolution experiment | STE4-WHI2::HphMX/ STE4-WHI2::KanMX, ade2–1, his3–11, leu2–3,112, trp1–1, ura3Δ::PFus1-yEVenus, bar1Δ::ADE2, hmlαΔ::LEU2, GPA1::NatMX |
| yGIL1556 - unlinked | MATa/a | Evolution experiment | WHI2::HphMX- STE4/whi2C85T::KanMX-STE4, ade2–1, his3–11, leu2–3,112, trp1–1, ura3Δ::PFus1-yEVenus, bar1Δ::ADE2, hmlαΔ::LEU2, GPA1::NatMX |
| yGIL1557 - linked | MATa/a | Evolution experiment | WHI2::HphMX-STE4/whi2C85T::KanMX-ste4G943T, ade2–1, his3–11, leu2–3,112, trp1–1, ura3Δ::PFus1-yEVenus, bar1Δ::ADE2, hmlαΔ::LEU2, GPA1::NatMX |

*Two alleles provided only for heterozygous loci in diploid genotypes.
†*ura3* alleles harbor spontaneous LOF mutations isolated from 5FOA.
‡ *ste4C958A* control allele to ensure there is no fitness effect of a synonymous PAM site substitution used in strain construction.
